# Visual context affects the perceived timing of tactile sensations elicited through intra-cortical microstimulation

**DOI:** 10.1101/2024.05.13.593529

**Authors:** Isabelle A. Rosenthal, Luke Bashford, David Bjånes, Kelsie Pejsa, Brian Lee, Charles Liu, Richard A. Andersen

## Abstract

Intra-cortical microstimulation (ICMS) is a technique to provide tactile sensations for a somatosensory brain-machine interface (BMI). A viable BMI must function within the rich, multisensory environment of the real world, but how ICMS is integrated with other sensory modalities is poorly understood. To investigate how ICMS percepts are integrated with visual information, ICMS and visual stimuli were delivered at varying times relative to one another. Both visual context and ICMS current amplitude were found to bias the qualitative experience of ICMS. In two tetraplegic participants, ICMS and visual stimuli were more likely to be experienced as occurring simultaneously when visual stimuli were more realistic, demonstrating an effect of visual context on the temporal binding window. The peak of the temporal binding window varied but was consistently offset from zero, suggesting that multisensory integration with ICMS can suffer from temporal misalignment. Recordings from primary somatosensory cortex (S1) during catch trials where visual stimuli were delivered without ICMS demonstrated that S1 represents visual information related to ICMS across visual contexts.

## Introduction

Tactile sensation is highly important for executing dexterous, adaptable movements (Ghez et al., 1995; Miall et al., 2021, 2019; Robles-De-La-Torre, 2006; Sainburg et al., 1995) and providing a sense of embodiment (Giummarra et al., 2008; Jeannerod, 2003; Tsakiris et al., 2010). In cases of spinal cord injury (SCI), motor and somatosensory abilities are impaired or fully lost below the level of the injury. Brain-machine interfaces (BMIs) provide a potential method to restore these abilities, by decoding motor intentions from neural activity (Collinger et al., 2013; Dekleva et al., 2021; Moses et al., 2021), and by using intra-cortical microstimulation (ICMS) in primary somatosensory cortex (S1) to elicit artificial tactile sensations (Armenta Salas et al., 2018; Flesher et al., 2016).

Motor BMIs have become more accurate and sophisticated over the last 15 years (Dekleva et al., 2021; Keshtkaran et al., 2022; Simeral et al., 2021; Willsey et al., 2022). In contrast, broadly viable somatosensory BMIs remain at the proof-of-concept stage (Flesher et al., 2021), although some principles mapping the relationship between ICMS and sensations have emerged. A higher ICMS current amplitude elicits sensations more often than a lower current amplitude, and the sensations tend to be rated as more intense (Armenta Salas et al., 2018; Bjånes et al., 2022; Flesher et al., 2016; Hughes et al., 2021). The perceived location of elicited sensations reflects the topographic organization of S1 according to where the stimulation microelectrode arrays are implanted (Armenta Salas et al., 2018; Flesher et al., 2016). However, it remains poorly understood how to achieve reliable, replicable sensations with controllable qualia because experiences of ICMS can vary widely across electrodes, participants, and experiments, even when stimulation parameters are kept constant (Armenta Salas et al., 2018; Callier et al., 2020; Flesher et al., 2016).

A somatosensory BMI implemented in the real world will necessitate ICMS being processed by the brain as part of a complex multisensory environment (Risso and Valle, 2022). To this end, understanding how ICMS is combined with other sensory inputs to produce perceptual experiences is essential. Reaction time studies have shown that artificial tactile sensations can be slower compared to real tactile inputs or visual stimuli (Bjånes et al., 2022; Caldwell et al., 2019; Christie et al., 2022; Godlove et al., 2014). However, while relative processing speeds between sensory modalities have been investigated, the dynamics of how they are integrated together are still unclear.

Given that visual and tactile stimuli are often paired together in the real world, the characteristics of the temporal binding window, or the period of time in which two stimuli are perceived as occurring simultaneously, needs to be mapped out with respect to ICMS and visual stimuli. It has been shown that the optimal timing needed to perceive peripheral nerve stimulation and visual stimuli as simultaneous is not always the same; peripheral stimulation in the leg must occur earlier than in the hand relative to visual stimuli to achieve optimal synchronicity (Christie et al., 2019b). Yet the temporal binding window between ICMS and vision remains unclear, and it is unknown what timings would be optimal to perceive ICMS and a visual cue as simultaneous.

In addition to timing considerations, it is also possible that vision can affect the qualia of ICMS-elicited sensations. Some work in lower limb amputees has shown that visual information can bias the localization of sensations elicited through peripheral nerve stimulation (Christie et al., 2019a). However, there has been little research on the potential effects of visual context on the neural processing and perceptual results of ICMS.

In this work, we explore the behavioral and neural results of pairing ICMS and visual stimuli in two tetraplegic patients implanted with microelectrode arrays in primary somatosensory cortex (S1) (**Fig. 1a**). In order to understand the importance of behaviorally relevant visual information to the perception of ICMS sensations, two visual conditions are used which differ in their level of realism (**Fig. 1c**). The visual stimuli are presented at varying temporal offsets relative to ICMS in order to better characterize the temporal binding window between ICMS and vision (**Fig. 1b**), and S1 recordings are examined during catch trials to assess the neural response to visual information related to ICMS. We find evidence that both visual context and ICMs current amplitude are capable of biasing qualitative aspects of ICMS-elicited sensations, and that the temporal binding window changes based on the behavioral relevance of visual context. Additionally, the point of peak simultaneity (PSS) of ICMS and vision varied substantially between the two participants, but was offset from zero in both, indicating an imperfect temporal alignment between ICMS and visual stimuli.

**Figure 1:**
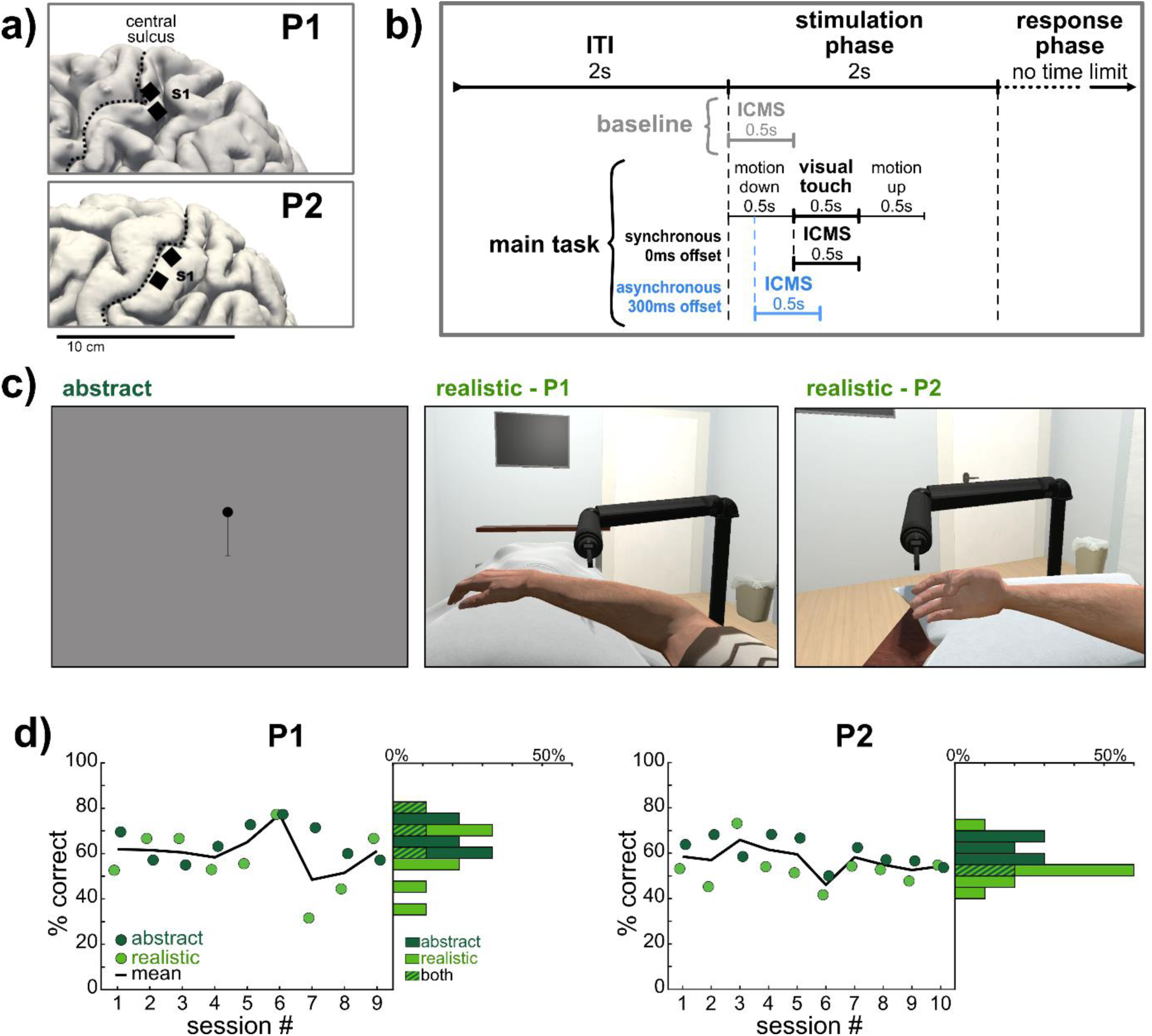
Experimental Methods and Paradigm. **a)** Microelectrode array implant locations (n=2), visualized using MRI on the cortical surface of each participant’s left hemisphere. **b)** Task time course. In the baseline, only ICMS was delivered. In the main task (either abstract or realistic), a visual cue was temporally linked to the ICMS at a given offset ranging between -300 and 300ms); 300ms offset is depicted. **c)** Sample frames from visual cues. In the abstract condition, the dot moved down to contact the end of the line. In the realistic condition, the robotic arm moved down to contact the virtual body: either the forearm (P1) or the index finger (P2). See **Supplementary Videos 1, 2, 3**. **d)** Scatter plot depicts behavioral accuracy across experimental sessions within each participant, quantified as the percentage of the trials where the participant reported a sensation, in which the participant’s reported order of stimuli (either vision first, ICMS first, or simultaneous) matched the ground truth stimulus order. Histograms to the right of the scatter indicate behavioral accuracy binned across sessions. The average number of trials felt per session in realistic runs was 18.2 (std=1.2) for P1 and 39.7 (std=4.1) for P2; in abstract runs the average was 19.7 (std=3.2) for P1 and 41.6 (std=5.8) for P2. Black line = mean across abstract and realistic runs.

Finally, we show that S1 represents information from visual stimuli relevant to ICMS in a context-independent manner. This study lays the groundwork for the implementation of BMIs using ICMS to elicit naturalistic sensations which can be temporally and perceptually integrated with the real-world environment.

## Methods

### Participants

As part of a brain-machine interface (BMI) clinical trial involving intracortical recording and stimulation, two tetraplegic participants (both male, age 33 & 39) with C5-level spinal cord injuries were recruited and consented. The microelectrode array implants of participant 1 (P1) were placed in three locations in the left hemisphere: the supra-marginal gyrus (SMG), ventral premotor cortex (PMv), and primary somatosensory cortex (S1). At the first experimental session, P1 was 5.5 years post-implant and 7 years post-injury. Participant 2 (P2) was implanted with microelectrode arrays in five locations on the left hemisphere: SMG, PMv, primary motor cortex (M1), anterior intraparietal area (AIP), and S1. At the first experimental session, P2 was 1 year post-implant and 3.5 years post-injury.

In total, two S1 arrays were implanted in each participant (**Fig 1a**). These arrays were 1.5mm SIROF-tipped (sputtered iridium oxide film) microelectrode arrays (Blackrock Neurotech, Salt Lake City, UT). P1’s arrays were 48-channel; P2’s arrays were 64-channel. Given the constraints of implanting arrays on the surface of cortex and the anatomy of S1, it is likely the S1 microelectrode arrays in both participants are located in Brodmann area 1 (BA 1) (Pandarinath and Bensmaia, 2022). Implant locations were determined based on fMRI motor and sensory localizer tasks. Additional information on the surgical methodology is available in (Armenta Salas et al., 2018).

All procedures were approved by the Institutional Review Boards (IRB) of the California Institute of Technology, University of Southern California, and Rancho Los Amigos National Rehabilitation Hospital.

### Experimental Paradigm

Three task conditions were tested in block format: a baseline, the realistic condition, and the abstract condition (**Fig. 1b**). During all runs, visual stimuli were shown to the participants using a Vive Pro Eye virtual reality headset (HTC Corporation, Taoyuan City, Taiwan) which was programmed using Unity. Nine sets of all three conditions were collected for P1; ten sets of just the realistic and abstract condition were collected for P2. For each participant, the same electrode was used to stimulate across conditions, which was selected to be highly reliable for eliciting tactile percepts based on earlier ICMS experiments.

In the baseline condition (18 trials per set), which always occurred before the other two conditions, P1 viewed grids of the upper body and hand on a gray background which remained static throughout the task. Each trial contained a 2s ITI, a 2s stimulation phase, and a response phase without a time limit. In each trial, 0.5s of ICMS was delivered immediately upon entering the stimulation phase, at one of three current amplitudes (30, 60, 100 µAmps) which were evenly sampled (5 trials each). All stimulation occurred on the same single channel, at 300 Hz, with a pulse-width of 200µs and an interphase of 60µs. The baseline also contained 3 catch trials where no stimulation was delivered. During the response phase, the participant was given an auditory cue (a beep) to verbally indicate whether or not he experienced a tactile sensation from ICMS. In the affirmative case he used the grids as references to indicate the sensation location. He also relayed the duration (‘Short’, ‘Medium’, or ‘Long’), a qualitative descriptor of the sensation (free word choice), and the intensity (on a subjective scale).

In the realistic and abstract conditions (P1: 40 trials per set within each condition; P2: 64 trials), the experimental phases and ICMS parameters were identical to the baseline, except a visual component was now included (**Fig. 1c**). In the abstract condition, the participants viewed a 2D black dot positioned at the top of a black line on a gray background (**Supplementary Video 1**). In the realistic condition, the VR headset was used to give the participants a first-person perspective of a body with a size, gender, and posture reflecting their own body (**Supplementary Videos 2, 3**), which was taken from the Microsoft Rocketbox Avatar Library (Gonzalez-Franco et al., 2020) (https://github.com/microsoft/Microsoft-Rocketbox/). In the VR environment, a virtual robotic, articulated arm with a narrow rod protruding from the end was positioned over P1’s virtual arm or P2’s virtual finger. P1’s forearm and P2’s index finger were selected as targets for the visually depicted touch to match the respective projected fields of the stimulated electrodes for each participant.

During the stimulation phase, both a visual and ICMS cue (same parameters as in the baseline) were delivered. In the realistic condition, the visual cue was the robotic arm performing a single tap of P1’s virtual arm or P2’s virtual finger, and in the abstract condition it was the dot moving along the line to tap the base of the line (**Fig 1c**, **Supplementary Videos 1, 2, 3**). In both conditions, the virtual cue was composed of 0.5s motion downwards, 0.5s contact (“visual touch”), and 0.5s motion upwards to the original position (**Fig. 1b**), such that the direction and magnitude of movement in the visual field were held constant across conditions.

The visual point of contact either to the virtual first-person body or to the end of the abstract line was depicted at varying times relative to the ICMS cue (P1: -300, -150, 0, 150, 300ms; P2: -300, -225, -150, -75, 0, 75, 150, 225, 300ms). In 12 of the trials in a set (P1: 40 trials total; P2: 64 trials total), they were presented simultaneously (0ms). Visual contact occurred before ICMS began in 12 trials for P1 and 24 trials for P2 (6 trials per individual offset). Similarly, visual contact occurred after ICMS began in 12 trials for P1 and 24 trials for P2. There were also 4 catch trials where the visual cue was delivered without ICMS. ICMS amplitudes (30, 60, 100µA) were sampled evenly within timing conditions. Within conditions, trials were pseudo-randomly shuffled. The order of the realistic and abstract condition blocks was alternated across days, while ICMS amplitudes were shuffled randomly within blocks.

During each trial in the realistic and abstract conditions, P1 reported if a tactile sensation was perceived, and if it was, he reported the perceived order of stimuli (‘vision first’, ‘ICMS first’, or ‘simultaneous’) as well as the location, intensity, duration, and qualitative nature of the sensation as in the baseline condition. During the realistic and abstract conditions, P2 only reported if a tactile sensation was perceived and the perceived order, due to time constraints during data collection.

### Data Collection

In total, nine sets of conditions were collected with P1 on nine unique days over 6 months. Ten sets were collected with P2 on six unique days over 3 months. Neural data was recorded from the S1 microelectrode arrays using a Neural Biopotential Signal Processor as 30,000 Hz broadband signals, and a CereStim96 device was used to deliver ICMS in S1 (Blackrock Neurotech, Salt Lake City, UT).

A central computer used custom MATLAB (MathWorks, Natick, MA) code with synchronized ICMS and visual outputs, the latter of which were displayed with a virtual reality headset (Vive Pro Eye, HTC Corporation, Taoyuan City, Taiwan). Eyetracking data was collected in both participants using the built-in camera and software in the VR headset, as well as custom Unity code.

Latencies to stimulus delivery were calculated and compensated for, resulting in a negligibly small unintended temporal offset of ICMS occurring an average of 5ms (std = 2ms) earlier than planned in P1 runs, and 10ms (std = 6ms) in P2 runs, relative to visual outputs across sessions.

### Data Preprocessing and Analysis

All analyses were performed using MATLAB R2019b (MathWorks, Natick, MA) unless otherwise noted. Data from P1’s S1 arrays were passed through a 180Hz notch filter in order to remove an electrical artifact which occurred throughout all recording sessions. Similarly, data from P2’s S1 arrays were passed through a 60Hz notch filter and a 920Hz notch filter in order to remove electrical artifacts which also occurred throughout all recording sessions. Multi-unit firing rates were computed from each channel’s broadband signals in 50ms bins without spike sorting (Christie et al., 2015; Dai et al., 2019), with a threshold of -3.5 times the noise RMS of the continuous signal voltage. These firing rates were aligned within each trial to the ICMS and visual stimuli presented. Firing rates were normalized within each run and each channel by calculating the mean baseline firing rate across the entire 2s ITI period, and dividing all firing rates in the session by this value.

Throughout the analysis, when multiple comparisons were performed, Bonferroni-Holm correction was performed to correct the p-values.

### Gaussian curve fitting

The percentages of trials where the participant reported simultaneous ICMS and visual percepts were fit to Gaussian curves (**Fig. 4**). Gaussians were fit to the raw percentages for each session and current amplitude reported in **Fig. 4a** and **c**, using MATLAB’s *fit* function, and restricted to peaks bounded by [0, 100] since the physical limits of simultaneous reports are 0% and 100%. A parametric bootstrap with 5000 iterations was used to assess variance (Christie et al., 2022). In the bootstrap, a binomial distribution *B(n,p)* was fit to the raw data at every time point, in which *n*=the number of trials at that time point and *p*=the percentage of trials that were reported as simultaneous. On every iteration of the bootstrap, these binomial distributions were sampled using MATLAB’s *binornd* function and Gaussian curves were fit to the resulting synthetic data. 95% confidence intervals on the Gaussians and their peaks were computed by examining the distribution of Gaussians generated over the bootstrap. The point of subjective simultaneity was taken as the peak of the fitted Gaussians. The just-noticeable difference (JND) was taken as the time in milliseconds between the peak and the 25% point of the Gaussian curve.

A Gaussian model was chosen for its simplicity, given the relatively sparse sampling of temporal offsets in the data. A more complex model runs the risk of overfitting to the data, and would require a larger number of time samples to better characterize the shape of the temporal binding window.

### Tuning Analysis

The tuning of multi-unit activity in each visual condition was assessed using the catch trials (P1: n=36; P2: n=40) collected during the realistic and abstract conditions, in which the visual cue was presented and the participant expected an ICMS-elicited sensation, but no ICMS was delivered. Tuning was computed via linear regression analysis in 100ms bins. In each time bin, normalized firing rates were compared to baseline firing rates, which were computed as the mean firing rates in the ITI, 1750ms to 750ms before the onset of the “stimulation” phase (**Fig. 1b**). Data for each channel was fit to a linear regression model based on the equation:

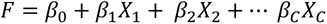

where *F* = vector of firing rates on each trial, *X* = one-hot-encoded matrix of time bin identity for each trial, *β* = estimated regression coefficients indicating level of tuning in each time bin, and *C* = number of time bins tested. *F* also included *n* additional entries corresponding to *β_0_* (*n* = the total number of catch trials), which contained the baseline firing rate calculated as indicated above. The null hypothesis β = 0 was assessed using a student’s t test within each channel and time bin, and if the null hypothesis was rejected then the channel was determined to be tuned to the visual information in comparison to the baseline firing rate. Within each channel, P values were corrected for multiple comparisons across time bins using the Bonferroni-Holm method.

To assess significant differences in numbers of tuned channels across time bins and conditions, a bootstrap analysis was run for 1000 iterations. In each iteration, the catch trials were randomly sampled with replacement and reassessed for tuning in each channel and time bin (**Fig. 5a**).

### RSA and MDS

Normalized firing rate data, binned in 0.5s phases, was examined using Representational Similarity Analysis (RSA, **Fig. 5d**) (Kriegeskorte, 2008; Nili et al., 2014). The python package *rsatoolbox* (https://github.com/rsagroup/rsatoolbox) was used to compute representational dissimilarity matrices (RDMs). The measure of dissimilarity used was cross-validated Mahalanobis distance with multivariate noise normalization (Walther et al., 2016), in which the noise covariance matrix is estimated and regularized towards a diagonal matrix to ensure that it is invertible. The cross-validated Mahalanobis distance is an unbiased measure of square Mahalanobis distance which also has a meaningful zero-point (Diedrichsen et al., 2021; Walther et al., 2016). A distance of zero between two conditions indicates the underlying neural data is fully indiscriminable, and the larger the Mahalanobis distance, the more these neural patterns are discriminable.

Data was cross-validated in session-wise splits (P1: 9 sessions; P2: 10 sessions), each containing the 4 catch trials collected per run for each condition and divided into 0.5s bins (‘ITI’, ‘Down’, ‘Touch’, ‘Up’). ‘ITI’ was composed of the last 0.5s of the ITI before the ‘Down’ phase began. The RDM generated from this data is symmetric across the diagonal, with meaningless zeros on the diagonal itself (**Fig. 5d**).

To better visualize the relationships in the RDM, multi-dimensional scaling (MDS) was applied using the MATLAB toolbox *rsatoolbox* (**Fig. 5e**, https://github.com/rsagroup/rsatoolbox_matlab)(Nili et al., 2014). MDS allows for distances in RDMs to be mapped to the 2D plane as faithfully as possible, using a metric stress criterion to arrange points without any assumptions of category structure. The stress between points is visualized with grey lines between points, stretched like rubber bands – the thinner the band, the more the true distances between points should be closer together to be fully accurate to the original high dimensional RDM.

### Eyetracking

Eye position (3D coordinates within the VR environment) and gaze direction (3D ray vector) within the VR environment were collected in both participants every 20ms, giving 6 features to describe eye movements which were averaged into 50ms bins to match the neural firing rate data. The headset software also returned a Boolean value indicating if the eye features were valid at each time point. If any timepoints within a 50ms bin were marked as invalid, the entire time bin was marked invalid. Eye movements were analyzed within catch trials to match the neural data analysis described above. Trials were used if at least 60% of time bins were marked as valid. In P1 (36 catch trials per condition), 12 realistic and 36 abstract trials were used. In P2 (40 catch trials per condition), 26 realistic and 30 abstract trials were used.

Within the valid trials, eye movement features were each normalized within each run by calculating the mean baseline firing rate across the entire 2s ITI period, and dividing all data in the session by this value (**Fig. S1a**). “Tuning” of eye features (**Fig. S1b**), RSA (**Fig. S1c**), and MDS (**Fig. S1d**) were then computed in the same manner as the neural data (see above), with the caveat that because the sets were missing trials, RSA was performed without cross-validation.

## Results

To understand the relationship between ICMS and visual context, behavioral and neural responses were recorded from two human tetraplegic patients (P1, P2) implanted with microelectrode arrays in S1 (**Fig. 1a**) (Blackrock Neurotech, Salt Lake City, Utah) as ICMS was delivered.

Three conditions were collected and examined with participant P1: the baseline, a realistic condition, and an abstract condition (**Fig. 1b, c**). During the baseline task, P1 reported when a sensation was elicited, that sensation’s anatomical location and intensity, and a one-word self-generated descriptor for the sensation’s qualitative nature. During the main task, P1 reported the same information, but also the relative order of ICMS and the visual cue, which was either depicted as an abstract or realistic touch (**Fig. 1c**, **Supplementary Videos 1, 2**). The visual cues were delivered at varying offsets relative to ICMS (-300, -150, 0, 150, 300ms). In all conditions, P1 only reported sensations localized to the right arm, between the wrist and the shoulder.

With participant P2, only the realistic and abstract conditions were collected and analyzed. Additionally, P2 only reported if a sensation was perceived, and the relative order of ICMS and the visual cue (**Fig. 1c**, **Supplementary Videos 1, 3**). Although P2 did not report sensation location on each trial, he reported after each study session that the sensations were localized on the tip of his index finger.

This set of experiments was designed to assess the effect of visual context on ICMS percepts, with particular interest in how these two stimuli would be percieved in temporal relation to one another. Additionally, catch trials without ICMS made it possible to examine the neural response of S1 to visual information.

Behavioral accuracy was computed by comparing the participants’ assessments of relative ICMS and visual stimulus order with the ground truth. There was no learning effect: accuracy did not meaningfully change over the different session days (**Fig 1d**, F-test vs constant model, P1: p=0.58; P2: p=0.14). For P1, behavioral performance was unaffected by the visual condition (**Fig 1d**, logistic regression test p=0.34), while P2’s performance was slightly better (p=0.01) in the abstract condition (mean = 60.5%, std = 6.3%) than in the realistic condition (mean = 52.8%, std = 8.4%).

### ICMS-elicited tactile sensations

On each trial, the participants reported whether or not they sensed a tactile percept. Three different ICMS current amplitudes were tested (100, 60, 30µA), and catch trials were also collected where no ICMS was delivered. P1 never reported a sensation in a catch trial. P2 reported one sensation during a catch trial; all data collected on that day was discarded and was not used for further analyses. There was a strong effect of ICMS current amplitude on the probability of P1 reporting a percept on a trial in the baseline trials (logistic regression test, p=1.7x10^-9^) as well as on the probability of both participants reporting a percept in both visual conditions of the main task (logistic regression test, P1: p=3.2x10^-33^; P2: p=1.9x10^-71^; **Fig. 2**). In both participants across conditions, at 100µA current amplitude, sensation detection was essentially at ceiling, and at 30µA, sensation detection was near floor.

**Figure 2:**
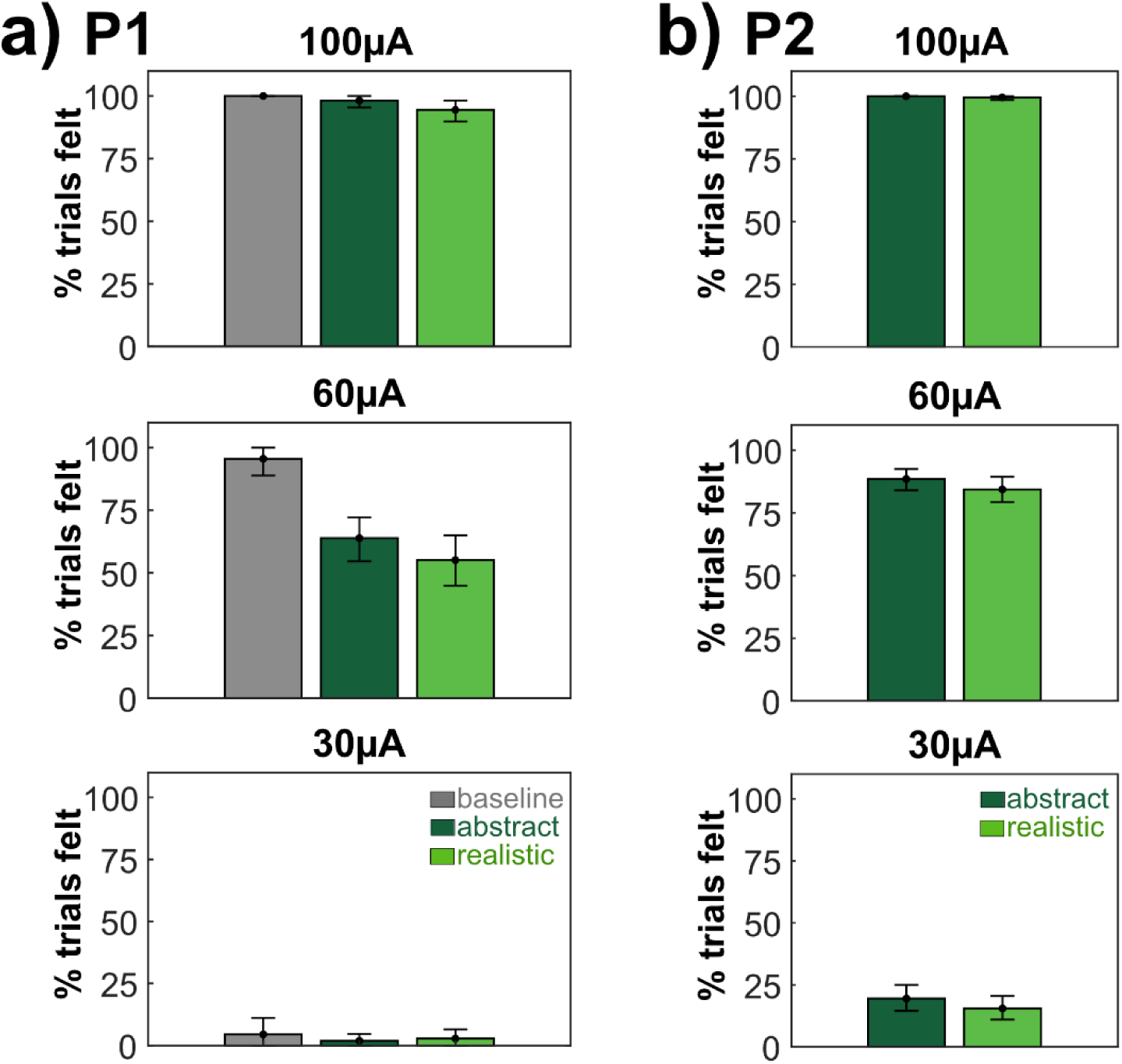
Tactile sensation rates across ICMS amplitudes. **a)** Percentage of trials eliciting a sensation across conditions and timing offsets in participant P1, sorted by ICMS current. Error bars represent 95% confidence intervals (CIs) assessed by bootstrapping values across 1000 iterations sampling from trials with replacement. **b)** Analysis identical to **(a)**, using data from participant P2.

In contrast, during 60µA ICMS trials, an effect of condition on the probability of reporting a sensation was evident (**Fig. 2**, P1: baseline mean=95.6%, 95%CI=[88.9,100]; abstract mean=63.9%, [54.6, 72.2]; realistic mean=55.1%, [44.9,65.0]; P2: abstract mean=88.5%, [84,92.5]; realistic mean=84.3%, [79.3,89.4]). The probability of a trial yielding a sensation was not different between realistic and abstract conditions (logistic regression test, P1: p=0.12, P2: p=0.19). In P1, the baseline elicited more sensations than either the realistic or abstract conditions (**Fig 2a**), which may be due to the lower cognitive load of the baseline compared to the main task.

When P1 reported a sensation, he also reported the intensity of the sensation using a subjective number scale (**Fig. 3a**) and a single word descriptor about how the sensation felt (**Fig. 3b, c**). Within realistic and abstract trials, there was no effect of condition or stimulus timing on intensity ratings, but there was an effect of ICMS current amplitude (3-way ANOVA, condition: p=0.69, timing: p =0.49, current: p = 1.5x10^-17^, all interaction effects p>0.05). Comparing 100µA and 60µA specifically, the greater amplitude led to greater intensity ratings (unpaired t-test, p=1.2x10^-21^).

**Figure 3:**
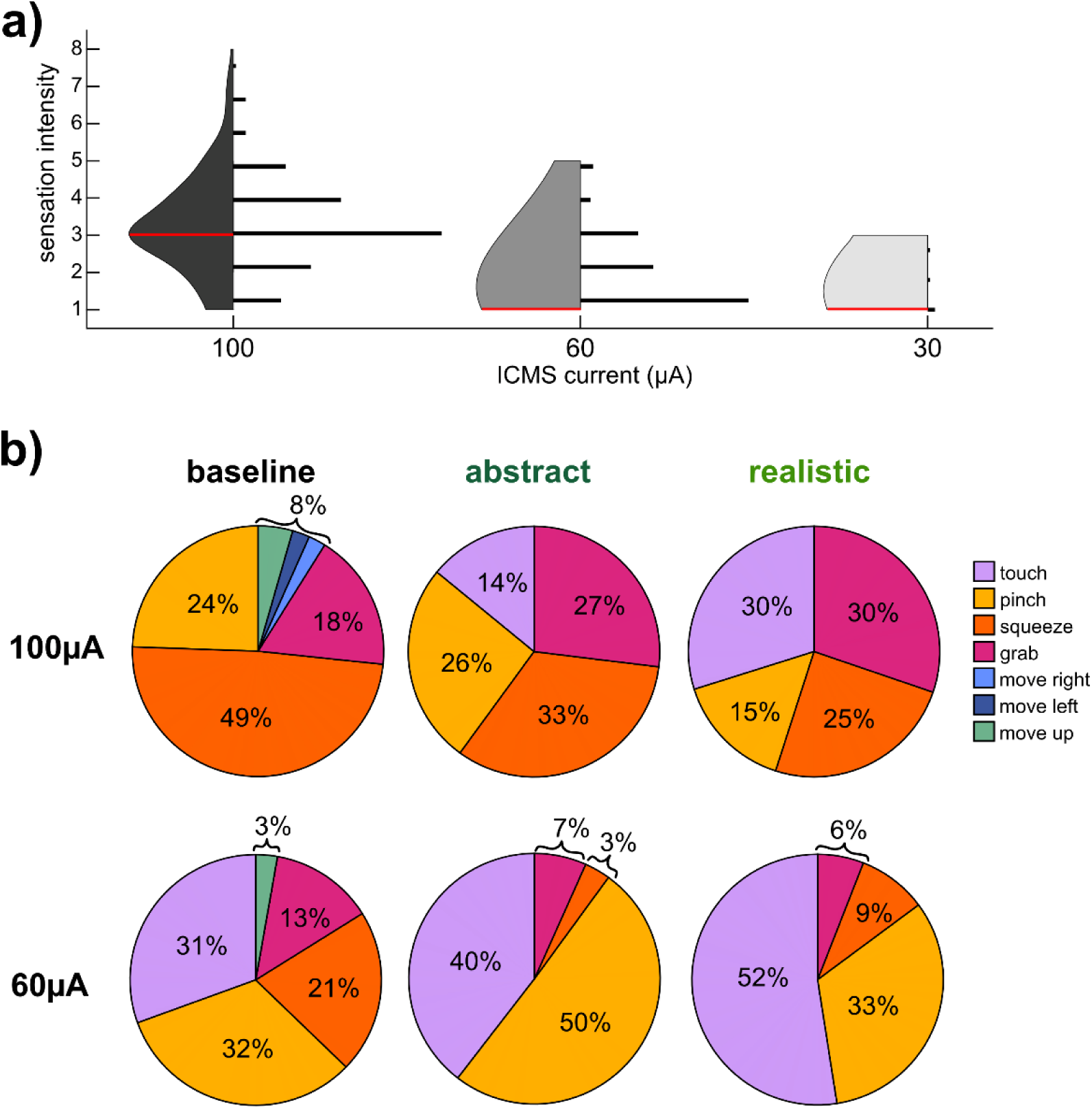
Qualia of ICMS-elicited tactile percepts in P1. **a)** Violin plot and histogram of reported sensation intensity separated by ICMS current and combined across abstract and realistic trials. Red line = median value. Histograms are to scale with the number of trials (n for 100µA = 83, 60µA = 67, 30µA =3 trials). **b)** Pie charts of sensation descriptors by condition and ICMS current amplitude. Only one word was used to describe each sensation. Top = 100µA (baseline n=45, abstract n=106, realistic n=102 trials). Bottom = 60µA (baseline n=43, abstract n=69, realistic n=59 trials).

Across 9 sessions, within 100µA trials, the word “touch” was used to describe the sensation in realistic trials a higher proportion of the time than in the baseline (**Fig. 3b**, 100µA trials Wilcoxon rank sum test, p=8x10^-4^), but the difference between abstract and baseline uses of “touch” was insignificant (p=0.24). Similarly, in 60µA trials, the highest proportion of “touch” sensations occurred in the realistic condition and the lowest occurred in the baseline, but this trend was not significant (**Fig. 3b**, 60µA trials realistic vs baseline: p=0.21, abstract vs baseline: p=0.21). Across all conditions, the word “touch” was used more in 60µA trials than 100µA trials (p=0.014).

### The temporal binding window between vision and ICMS

In every trial of the realistic and abstract conditions that the participants reported an ICMS-elicited sensation, they reported the perceived order of the ICMS and visual cue. Specifically, participants could either state that one stimulus came before the other or that they occurred simultaneously (**Fig. 4a, c**). Overall, the likelihood of P1 giving a “simultaneous” answer was affected both by timing offset (logistic regression test, p=0.01) and visual condition (p=0.001), but not current amplitude (p=0.27). The likelihood of P2 giving a simultaneous answer was affected by timing offset (p=4.2x10^-5^), visual condition (p=0.04), and current amplitude (p=7.7x10^-15^).

To further examine how visual condition affected perceived timing, participant responses were examined within the two current amplitudes tested with the largest number of perceived sensations: 100µA and 60µA (**Fig. 2**). Unsurprisingly, in both conditions and at both current amplitudes, the participants detected the correct order most easily in the -300 and 300ms offset trials (**Fig 4a, c**). In P1, within each ICMS current amplitude, the areas under the “simultaneous” curves were different between realistic and abstract conditions (100µA: p=3x10^-4^; 60µA: p=0.005). In P2, the areas were only different within 100µA trials (100µA: p=0.04; 60µA: p=0.11).

To better quantify when stimuli were perceived as occurring synchronously within 100µA trials, the “simultaneous” curves (**Fig. 4a, c**, black lines) were fit to Gaussians which allowed for interpolation between the tested timing offsets (**Fig. 4b, d**). The variability in the data was assessed using a parametric bootstrap (see **Methods**)(Christie et al., 2022). The point of subjective simultaneity (PSS), defined as the peak of the fitted Gaussians, occurred when ICMS preceded the visual cue for both visual conditions in P1 (abstract: 72.2ms, 95% CI = [47.0, 94.2]; realistic: 124.2ms [86.8, 166.8]). In P2, the opposite was true: the PSS occurred when the visual cue preceded ICMS (abstract: -11.4ms, 95% CI = [-38.7, 18.6]; realistic: -63.2ms, 95% CI = [-95.9, -30.7]).

**Figure 4:**
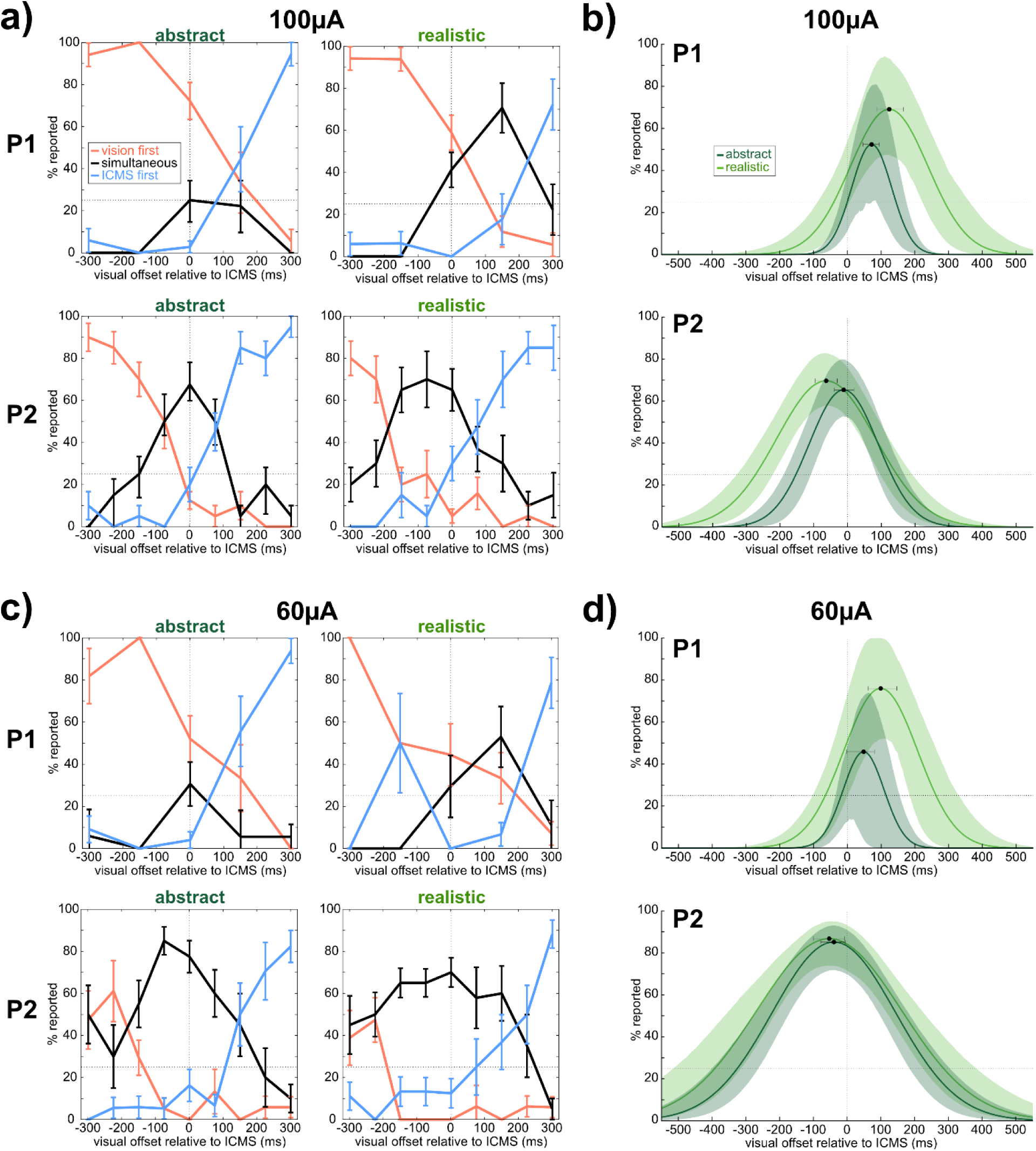
The temporal binding window between vision and ICMS. **a)** Reports of stimulus order relative to the ground truth amongst 100µAmp trials in each participant (top row: P1; bottom row: P2). A negative timing offset indicates the visual stimulus preceded ICMS. Horizontal dotted line indicates 25% mark. Error bars represent SEM. **b)** Gaussian curves fit to the “simultaneous” points (black lines) in **(a)** for each participant. Black dots indicate curve peaks. Shaded area and error bars on peaks represent 95% CIs generated through a parametric bootstrap fit to 5000 synthetic versions of the data modeled with a binomial distribution (see Methods). **c)** Analysis identical to **(a),** in 60µAmp trials. **d)** Analysis identical to **(b)**, in 60µAmp trials.

The just-noticeable difference (JND), defined as the time in milliseconds between the peak and the 25% point of the Gaussians, was larger in realistic trials (P1: 164.0ms [113.6, 221.8]; P2: 209.6ms [167.7, 247.4]) than in abstract trials (P1: 72.3ms [24.0, 95.6]; P2: 141.7ms [105.1 169.8]), although this difference was only significant in P1. In P1, the 25% point on the left side of the Gaussian was not different between realistic and abstract conditions, whereas the 25% point on the right side was different across conditions: the realistic condition had a larger offset than the abstract condition. In P2, the reverse was true: the 25% point on the left side was larger in the realistic condition than in the abstract condition, while the right-hand 25% point was not different across conditions.

The data within 60µA trials were also fit to Gaussians using the same procedure (**Fig. 4d**). In P1, the 60µA realistic PSS (98.8ms [61.3 146.5]) was not significantly different from the abstract PSS (48.3ms [-1.1, 80.4]). The 25% points of the Gaussians were also not different from one another between the realistic condition and the abstract condition. However, in P1 the realistic JND was larger in the realistic condition (164.0ms [98.1, 217.2]) than in the abstract condition (66.6ms [0.1, 95.7]). In P2, the realistic PSS (-53.0ms [-100.0, -8.2]) was not significantly different from the abstract PSS (-39.1ms [-78.4, -5.9]). The JNDs were also similar across conditions in P2 (realistic: 338.7ms [281.3, 430.6]; abstract: 286.1ms [230.4, 339.1]). Neither the left nor the right 25% point of the Gaussian was different across conditions.

### S1 neural responses to visual stimuli

In both participants, neural activity was recorded in the S1 microelectrode arrays during catch trials, when no ICMS was delivered. Multi-unit channel firing rates were computed, and the tuning of channels relative to baseline while visual stimuli were delivered was assessed using a linear regression analysis (**Fig. 5a, b**). In both P1 and P2, the highest number of tuned channels in the realistic condition was in the -0.1s to 0s bin prior to visual touch, during end of the “motion down” phase. With respect to the abstract condition, the peak number of tuned channels occurred in the bin immediately after “visual touch” phase onset, 0 to 0.1s in both participants. The timecourse of two example tuned channels from each participant are shown in **Fig. 5b**.

**Figure 5:**
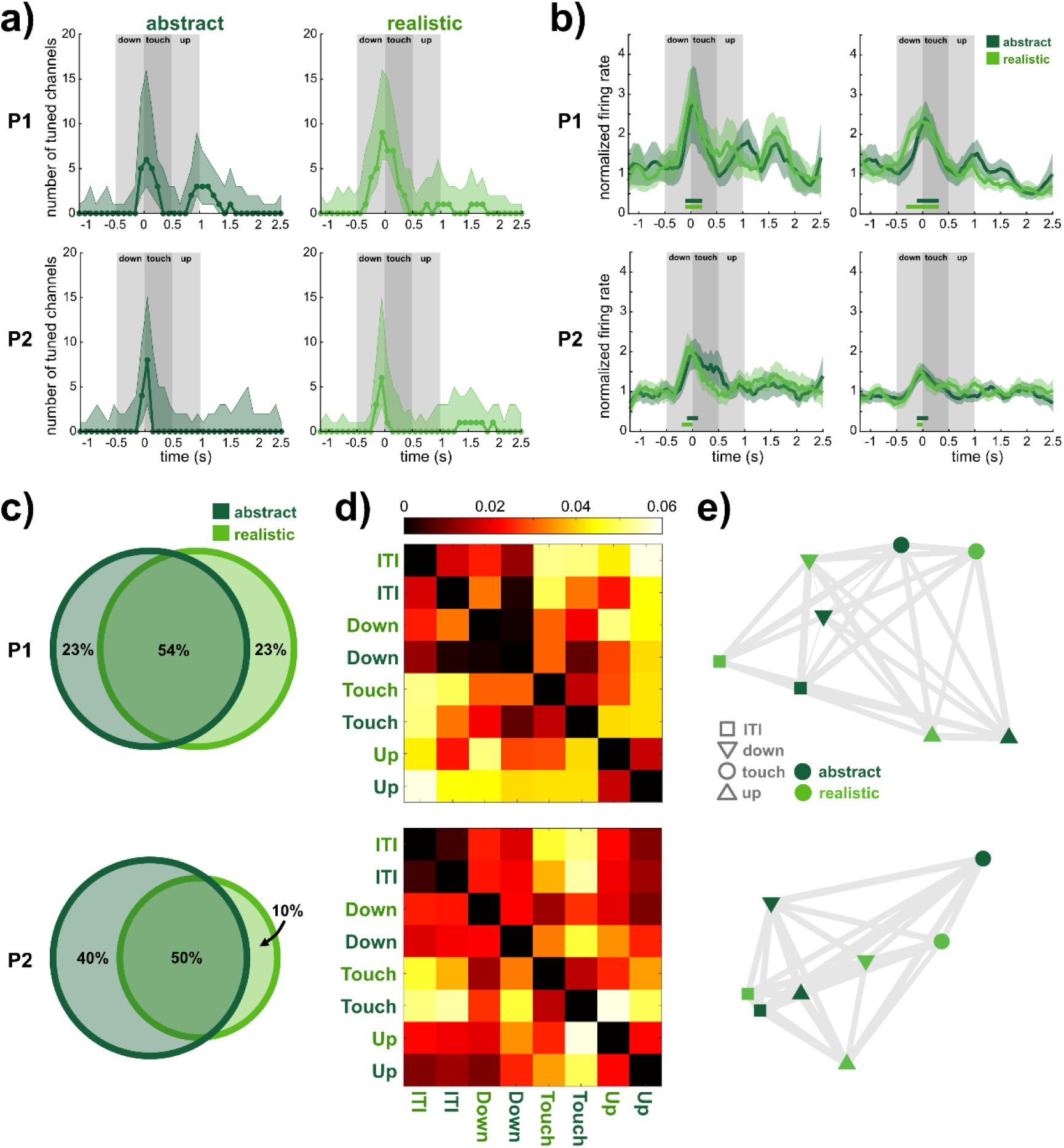
Neural activity during catch trials. **a)** Number of channels tuned during catch trials relative to baseline (P1 n=96 channels, P2 n=128), separated by condition. Time is aligned to the onset of the “touch” phase of the visual stimulus. Shaded area indicates 95% CIs computed by bootstrapping across trials (P1 n=36 trials, P2 n=40) over 1000 iterations. Tuning was assessed by linear regression analysis (see Methods). **b)** Example tuned channels with firing rates averaged across catch trials, by condition. For visualization only, firing rates were smoothed using a first order Savitzky-Golay filter. Horizontal bars indicate the times in which the channels were tuned relative to baseline, color-coded by condition. Shaded area corresponds to SEM. **c)** Venn diagrams depicting the overlap in channels tuned to abstract and realistic conditions. Percentages are based on the total number of channels tuned overall (P1: 13 channels, P2: 10 channels). **d)** Representational dissimilarity matrices (RDM) of multi- unit neural activity across all channels. Heatmap indicates distances between neural activity patterns associated with each condition and task phase (e.g. realistic ITI in top left), which are computed as the cross-validated Mahalanobis distance with multivariate noise correction; a distance of 0 indicated conditions are statistically indistinguishable. Each phase represents 0.5s of averaged firing rates; the ITI is based on the 0.5s immediately prior to “down” phase of visual stimulus. **e)** Multi-dimensional scaling (MDS) plots of the RDMs in **(d)**. Axes are arbitrary. Grey lines between icons are “rubber bands” whose thickness is based on the goodness of fit of the scaling (P1: Pearson’s r=0.93, p=1.5x10^-10^; P2: r=0.97, p=4.9x10^-17^). Thinner, more “stretched” bands indicate that the icons are closer together in the original high-dimensional space than they are shown to be.

In P1, 10 channels exhibited tuning during the realistic visual animation and 10 exhibited tuning during the abstract visual animation. In P2, 6 channels exhibited tuning during the realistic visual animation and 9 exhibited tuning during the abstract visual animation. This measure was calculated by counting the number of unique channels tuned in any of the 100ms bins during the entire visual animation including down, touch, and up (total animation length: 1.5s). The overlap in channels tuned to realistic and abstract conditions during the visual animation (1.5s total) was quantified (**Fig. 5c**). In both participants, more channels were tuned to both realistic and abstract conditions, than were tuned to either condition individually.

To understand the population response in S1 to the visual stimuli, Representational Similarity Analysis (RSA) was employed on the firing rates of all channels in each participant (**Fig. 5d**) (Kriegeskorte, 2008), using cross-validated Mahalanobis distance with multivariate noise correction (Walther et al., 2016). Multi-dimensional scaling (MDS) was used to visualize the computed distances between condition phases (**Fig. 5e**) (Nili et al., 2014). In P1, the data were grouped more tightly by task phase (ITI, down, touch, up) than by condition (abstract, realistic), as assessed by a Wilcoxon rank sum test on distances within phases/across conditions vs distances within conditions/across phases (p=0.03). Qualitatively, this is apparent in the MDS where icons are grouped by phase, but the conditions are intermixed together (**Fig. 5e**), which indicates that the neural activity was similar between the two visual conditions, and varied along the timecourse of the task in both conditions. In P2, a similar grouping is visually present to some extent (**Fig 5e**), but the distances within phases/across conditions were not significantly different compared to the distances within conditions/across phases (p=0.13).

It is possible that eye movements confounded the neural data, given the importance of visual information during this task. To assess this possibility, catch trials with valid eye tracking data were analyzed using the same linear regression analysis as the neural data (**Fig. S1b**; P1: 36 abstract trials, 12 realistic; P2: 30 abstract, 26 realistic). None of the 6 eye movement features were tuned relative to baseline during the vast majority of the visual animation in both participants, and the earliest that tuning occurred was immediately after the visual animation in P1, and during the final 0.25s of the 1.5s animation in P2 (**Fig. S1a, b**). The eye movement features were also used to perform RSA using Mahalanobis distance with multivariate noise correction (**Fig S1c**), and mapped to two dimensions using MDS (**Fig. S1d**). Eye movements were highly clustered according to visual condition compared to task phase (Wilcoxon rank sum test on distances within phases/across conditions vs distances within conditions/across phases, P1: p=0.001; P2: p=0.001) as is visually apparent in the MDS plots.

## Discussion

Behavioral and neural data from two tetraplegic participants (P1, P2) receiving ICMS in S1 (**Fig. 1a**) while observing visual abstract or realistic touch cues (**Fig. 1c, Supplemental Videos 1, 2, 3**) were examined. Visual cues were delivered at varying temporal offsets relative to ICMS (**Fig. 1b**) and the participants reported the perceived order of these cues. P1 also reported descriptive information about the ICMS-elicited tactile sensations. This experiment yielded three main findings: 1) the qualitative percepts of ICMS-elicited sensations are influenced by visual information and ICMS current amplitude; 2) the temporal binding window between ICMS and vision varies based on the behavioral relevance of the visual stimuli; 3) S1 represents visual information relevant to ICMS in a context-independent fashion.

### ICMS-elicited sensations are affected by visual information and ICMS current amplitude

This work replicates the previously known result that higher ICMS current amplitudes tend to elicit sensations more frequently (**Fig. 2**), and the sensations tend to be of higher intensity (**Fig. 3a**) (Armenta Salas et al., 2018; Flesher et al., 2016; Hughes et al., 2021). However, we also find an interaction between current amplitude and vision. During 60µA trials, the percentage of trials which elicited a sensation through ICMS were significantly higher in the baseline than in either of the visual conditions (**Fig. 2a**). Since 60µA ICMS is typically perceived to be lower intensity than 100µA ICMS (**Fig. 3a**), it is closer to the perceptual detection threshold. The added cognitive load of attending to a visual stimulus as well as attending to ICMS may be responsible for lower rates of reported sensations. Indeed, this behavioral result is predicted by the divisive normalization model of attention which suggests that when neurons respond strongly to attended stimuli, neural responses to other stimuli are proportionally suppressed (Lee and Maunsell, 2009; Reynolds and Heeger, 2009).

We also demonstrate an interaction effect of ICMS current and visual condition on the qualia of sensations elicited in P1: “touch” was used as a descriptor more often at 60µA than at 100µA, and within 100µA the word “touch” was used more often in the realistic condition than in the baseline (**Fig. 3b**). The increased use of the word “touch” in lower current amplitude trials may be because, within the participant’s subjective framework, the word “touch” could represent a tactile stimulus that is inherently of a lower intensity than other descriptors like “squeeze” or “grab.” Additionally, of the words the participant used to describe sensations, which were freely chosen by him, “touch” appears to be the one which most closely matches the visual touch depicted in the realistic condition.

Although previous work has shown that visual information can bias the perceived location of peripheral intraneural stimulation (Christie et al., 2019a), the effect reported here suggests that visual information can also bias the qualitative percept of ICMS-elicited sensations. When ICMS is employed on the same electrode, with the same parameters, in the same participant, widely varying qualia often result (**Fig. 3b**, baseline charts) (Armenta Salas et al., 2018; Flesher et al., 2016). A viable neural prosthetic would ideally be able to elicit naturalistic sensations of specific qualia as needed (Tabot et al., 2015). If visual information can stabilize ICMS percepts to some extent, this has the potential to be highly important to the development of such a tactile neural prosthetic.

### The behavioral relevance of visual stimuli influences the temporal binding window

Visual stimuli were presented at varying temporal offsets (0-300ms) relative to ICMS (**Fig. 4**). In P1, the PSS occurred when ICMS preceded the visual cue, with a lag of approximately 50-125ms depending on visual condition and ICMS current amplitude (**Fig. 4b, d**). In P2, the PSS occurred when the visual cue preceded ICMS, with a lag of approximately 10-65ms. The variance in the PSS of the two participants may be due to individual differences, differences between the properties of stimulating electrodes, or differences in microelectrode implant location – or some combination of these factors. We do not conclude that there is some constant temporal offset between ICMS and vision, but rather that future ICMS applications should not assume that the temporal binding window is centered at zero. This finding echoes prior work with amputees, which has shown that the PSS of peripheral nerve stimulation shifts depending on whether the upper or lower limb is stimulated (Christie et al., 2019b). While the stimulation delivered in this work is intra-cortical rather than peripheral, and both participant’s microelectrode arrays were implanted in putative S1, P1’s array is located in the arm region (Rosenthal et al., 2023) and P2’s array is located in the finger region (see **Methods**); this difference in cortical sub- regions may be sufficient to create processing time differences.

Aside from anatomical considerations, the sensory processing systems of different individuals are often equipped with different priors about the environment, biasing them towards one perceptual experience or another. A striking example of this phenomenon in the visual system is the viral photo of a dress that is perceived as either blue-black or white-gold due to varying assumptions by the visual system about the photo’s illumination (Lafer-Sousa et al., 2015). It is possible that P1 and P2 have different biases with respect to how they process visuo-tactile information.

Finally, stimulation occurred on a single electrode in each participant, chosen due to its reliability in eliciting tactile percepts in prior experiments. It is highly possible that multichannel stimulation would give rise to a different PSS than single channel stimulation, even within a single participant, given that reaction times to ICMS are shorter with multichannel stimulation than single channel stimulation (Bjånes et al., 2022). While outside of the scope of this work, our findings highlight the need to determine how the PSS changes across ICMS stimulation parameters and brain areas, and individual patients. Regardless of the direction of the offset, the existence of a perceptual lag between vision and ICMS in both participants indicates that the parameters used here, which are standard in the field, do not allow ICMS to be perfectly integrated into a temporally well-aligned multisensory experience.

Although the PSS remained relatively consistent within each participant across visual conditions, the width of the temporal binding window varied between visual conditions. In both participants, during 100µA trials, both the area under the curve of “simultaneous” answers (**Fig. 4a**) and the JND (**Fig. 4b**) were larger in realistic trials compared to abstract trials, indicating the temporal binding window between ICMS and vision is larger in the realistic condition. In other words, the participant was more likely to perceive ICMS and vision as occurring synchronously in the realistic condition, and more likely to assign an order to the stimuli in the abstract condition.

In 60µA trials, the area under the curve and the JND were larger in realistic trials for P1, just as in 100µA trials (**Fig. 4c, d**). In P2, the fitted Gaussian realistic curve was larger than the abstract one, and the realistic JND was larger than the abstract JND, although these differences were not significant. Since 60µA ICMS is less perceptible than 100µA ICMS (**Fig. 2**)(Armenta Salas et al., 2018; Flesher et al., 2016; Hughes et al., 2021), P2 may have integrated this weaker signal more easily with the visual cue across visual conditions, resulting in more “simultaneous” answers overall. Supporting this theory, we report an effect of current amplitude in the rate of “simultaneous” answers for P2, but no such effect for P1.

As a whole, the temporal binding window results indicate that a behaviorally relevant touch input allows the brain to more easily link visual and ICMS inputs together causally, and view them as happening as part of the same event, while in an abstract context, visual and ICMS inputs are more likely to be interpreted as two separate events. Since a somatosensory neural prosthetic using ICMS would be deployed in a real world environment, this result is encouraging because it supports the idea that the brain is able to combine multisensory realistic inputs with artificial stimulation to generate visually plausible sensations (Christie et al., 2019a).

It is possible that changes to the temporal binding window could be due to differences in task difficulty across the visual conditions. The abstract condition is a comparatively simple visual stimulus (**Fig. 1c, Supplementary Video 1**), centered in the visual field, so it may be easier to estimate timings in this condition than in the more complex realistic condition (**Fig. 1c, Supplementary Videos 2, 3**). Yet, the visual conditions were relatively well-matched in terms of difficulty: P1 performed equally well at assessing stimulus order between visual conditions, and although P2 performed slightly better at the abstract version of the task, the accuracy histograms of the two conditions were highly overlapping for both participants (**Fig. 1d**). Additionally, the ability of the participants to gauge the timing between ICMS and visual cues was not affected by learning, as across the experimental sessions (P1: n=9; P2: n=10), there was no change in the participants’ ability to accurately assess stimulus order. It is therefore unlikely that learning over time or task difficulty were major confounds in these findings.

### S1 represents ICMS-relevant visual content in a context-independent fashion

Examining catch trials during which visual stimuli were presented without ICMS, we find that a significant number of channels are tuned to visual touches relative to baseline activity in both participants (**Fig. 5a**). In particular, S1 activity is most tuned during the initial onset of the visual touch. Given that a total of 13 S1 channels in P1 and 10 S1 channels in P2 were tuned during the visual animations, it is clear that S1 reflects some component of the visual stimulus even when there is no ICMS or physical tactile event. While abstract trials elicited tuned activity peaking during the first 100ms of the visual touch onset, tuned neural activity during realistic trials peaked 100ms before the onset of visual touch, during the “motion down” period, indicating possible preparatory or predictive activity (Kimura, 2021).

There was substantial overlap between tuned responses to abstract and realistic trials in S1. Many of the channels tuned to the realistic condition were also tuned to the abstract condition (**Fig. 5b, c**). In a population-level analysis, RSA demonstrated that neural activity was not grouped by visual condition, as might be expected if the S1 response to realistic visual touches was drastically different from abstract visual touches, but instead neural activity was grouped by task phase (**Fig. 5d**). This grouping was significant in P1, but not for P2, although it was qualitatively present in the MDS for both participants **(Fig. 5e**). As discussed above, a possible reason for the variance between participants is that the locations of the microelectrode arrays are in different topographic regions of S1 (P1: arm area; P2: finger area), and may be positioned slightly differently within the tactile processing hierarchy (**Fig. 1a**).

Taken as a whole, these results indicate that S1 represents information contained in visual stimuli, and that this information generalizes across abstract and realistic conditions to some degree. Given that abstract and realistic stimuli are very visually distinct (**Fig. 1c, Supplementary Videos 1, 2, 3**), it is unlikely that S1 is representing the actual visual inputs themselves. In the lateral intraparietal area, a region that responds to passive visual stimuli and to saccade planning and execution, there is prior work demonstrating that the multisensory requirements of a task can cause a neural population to become tuned to auditory information in addition to visual information (Grunewald et al., 1999), and that these tuned responses are supramodal, linking oculomotor behavior to selected auditory targets (Linden et al., 1999). Similarly, in the experiments presented here, S1 is representing some aspect of the visual information that is relevant to the tactile aspect of the task and which generalizes across the two visual contexts in order to differentiate baseline and visual touch activity.

Eye movements represent a potential confound for these results – S1 could simply be reflecting changes in eye movement pattern during the different task phases. To control for this possibility, eye movements were recorded in both participants in catch trials, and no eye movement features were tuned relative to baseline during the first 1s of the visual animation (“down” and “touch” phases, **Fig. S1a, b**), in either visual condition. Unlike the neural data, eye movements were also overwhelming clustered according to visual condition in an RDM analysis (**Fig. S1c, d**). It is therefore highly unlikely that eye movements contribute to the neural patterns discussed here.

Prior work with one of the participants in this study (P1) showed that S1 did not respond to visually depicted touches without a physical tactile stimulus accompanying them (Rosenthal et al., 2023). In contrast, this study shows that S1 does reflect information in visual stimuli without any tactile stimuli or ICMS. This difference in results supports a hypothesis suggesting that task design has a large effect on whether S1 represents visual information related to touch (Dionne et al., 2013; Rosenthal et al., 2023). Rosenthal et al. (2023) used a passive design, in which the participant merely observed tactile stimuli, whereas this study implemented a more active task in which the participant was required to describe perceived tactile sensations and report the order of ICMS and visual stimuli. This effect of task design on S1 modulation by visual stimuli can also be seen in the neuroimaging literature. Experiments with an active task tend to find that S1 responds to observed touches (Blakemore et al., 2005; Bufalari et al., 2007; Ebisch et al., 2008; Kuehn et al., 2018, 2013; Longo et al., 2011; Schaefer et al., 2009), while experiments with a passive task, or a task that is not touch-related, tend to find the opposite (Chan and Baker, 2015; Keysers et al., 2004; Morrison et al., 2004). Visual information not related to the tactile stimulation also does not modulate somatosensory cortex (Espenhahn et al., 2020). This effect of passive verses active tasks has a parallel in non-human primates. The lateral intraparietal area typically responds to passive visual stimuli but not passive auditory stimuli. After training on a saccade task with auditory cues, neurons become responsive to auditory stimuli which represent saccade targets (Grunewald et al., 1999). It is therefore likely that sensory processing, from early cortical stages like S1 to posterior parietal cortex, can flexibly incorporate multisensory information in a context-dependent way.

Given that S1 can reflect visual stimuli based on task relevance, it is likely that behavioral relevance plays some role in what S1 represents in a given context (Chapman and Meftah, 2005; Dionne et al., 2013; Popovich and Staines, 2014). Such behavioral modulation may also underlie the visual enhancement of touch, a phenomenon in which tactile perception is improved when the body part being touched is visible, even if the visual input is non-informative about the touch (Colino et al., 2017; Haggard et al., 2007; Kennett et al., 2001; Press et al., 2004; Tipper et al., 2001). It may also play a role in S1’s ability to reflect top-down concepts like affective significance, motor planning, and imagined touches (Ariani et al., 2022; Bashford et al., 2021; Gale et al., 2021; Yoo et al., 2003).

However, it also seems apparent that when S1 represents information from vision, this encoding is temporally closely locked to the visual touch itself. In this work, we see that the onset of tuned responses never occurs more than 200ms ahead of visual touch onset, and the majority of responses occur with 100ms of visual touch onset, despite the visual animation starting 500ms before visual touch onset (**Fig. 5a**). In prior work with participant P1, visual information predictive of when a physical touch would occur did not activate S1 ahead of the physical touch (Rosenthal et al., 2023), and while the execution of a motor imagery task activates S1, visual cues relating to motor imagery are not sufficient to meaningfully activate S1 in the 4 seconds prior to performing the imagery (Jafari et al., 2020). Therefore while visual information is capable of modulating S1 (Rosenthal et al., 2023), there is a limited predictive effect of visual information in S1. These results are indicative of higher order efference copy signals that is computed outside of S1 as a component of a forward model, perhaps to be used for predicting the sensory consequences of a movement. Similarly, the visual response documented here is likely implemented by higher order brain areas which represent the task requirements and compute a relevance threshold for different sensory inputs which can then be implemented in modality specific early processing areas like S1.

Finally, the fact that the results in this work are consistent with other characterizations of S1 in tactile studies indicates that S1 processes visual stimuli related to ICMS in some similar ways to visual stimuli related to real physical touches. This suggests that ICMS can be a valid substitute for physical touches in tactile tasks, not only in terms of behavioral performance (Berg et al., 2013; Klaes et al., 2014; Tabot et al., 2015), but also in terms of how the stimulation is processed within the somatosensory system.

## Conclusion

In order to understand the behavioral and neural relationship between ICMS and vision, we examined responses to paired visual and ICMS stimuli at varying temporal offsets in two tetraplegic patients. This dataset yielded two behavioral findings. The first is that the interpretation of ICMS-elicited sensations is affected by ICMS current amplitude, as well as by visual content. The second is that the temporal binding window between ICMS and vision varies in offset but is not necessarily centered at zero, and that the size of the temporal binding window is affected by the behavioral relevance of the visual stimulus. Studies of ICMS frequently examine elicited sensations without including any other type of sensory context (Armenta Salas et al., 2018; Flesher et al., 2016; Hughes et al., 2022, 2021). While these studies represent important foundational work, it will be important for a real-world BCI to fully understand how ICMS interacts with a richly complex sensory environment in order to stabilize touch percepts and create temporally aligned, unified multisensory experiences.

By examining the neural encoding of catch trials in which visual touches were present without ICMS, this work also adds to our understanding of how S1 represents visual information related to tactile sensations. We find that in an active task, S1 firing rates change during a visual touch relative to baseline, in a relatively constant way across visual contexts. This finding supports the idea that high level task-related variables in visual stimuli are represented in S1 and modulated by higher order cognitive brain areas based on attention.

Further experiments should aim to investigate these findings across a larger set of ICMS parameters, to better understand how the temporal binding window can vary based on stimulation properties. Additionally, it will be important to explore in more depth how ICMS is integrated with other sensory systems and with different environmental contexts, as well as how S1 processes these inputs. By understanding multisensory ICMS integration both behaviorally and neurally, we can better use ICMS to design stable, naturalistic artificial tactile sensations.

## Limitations of the Study

While this experiment expands our understanding of how vision is integrated with ICMS, it is limited in scope. Testing two participants allows for an initial understanding of how individual differences contribute to our results, but further work will be necessary to fully characterize the possible variance across patients. Additionally, a limited set of ICMS parameters was examined, and it is likely the precise temporal relationship between ICMS and vision will shift if different parameters are tested. Finally, it is possible that demand characteristics of the task influenced P1’s description of sensation qualia, although we note that P1 never reported a sensation in a catch trial, and reported significantly fewer sensations at lower current amplitudes, despite never being informed of the current amplitude being delivered on any given trial.

## Acknowledgements

We thank the participants of this study for their effort and dedication to the study, and S. Wandelt and W. Griggs for helpful discussions and insights.

## Author Contributions

I.A.R., L.B., D.B., and R.A.A. designed the study. I.A.R. developed the experimental tasks. I.A.R., L.B., and D.B. collected data. I.A.R. analyzed the results. I.A.R., L.B., and D.B. interpreted the results. I.A.R. wrote the paper. I.A.R., L.B., D. B., and R.A.A. reviewed and edited the paper. K.P. coordinated regulatory requirements of clinical trials. C.L. and B.L. performed the surgery to implant the microelectrode arrays.

## Supplementary video titles and legends

**Supplementary video 1:** *Sample abstract visual stimulus.* Related to **Figure 1c**. Stimulus was presented using a virtual reality headset. The virtual reality environment was constructed to mimic the room in which the experiment was performed.

**Supplementary video 2:** *Sample realistic visual stimulus for P1*. Related to **Figure 1c**. Stimulus was presented using a virtual reality headset. The virtual reality environment was constructed to mimic the room in which the experiment was performed, and the human arm and hand were chosen to be visually similar to the participant’s own body.

**Supplementary video 3:** *Sample realistic visual stimulus for P2*. Related to **Figure 1c**. Stimulus was presented using a virtual reality headset. The virtual reality environment was constructed to mimic the room in which the experiment was performed, and the human arm and hand were chosen to be visually similar to the participant’s own body.

---

**Figure S1:**
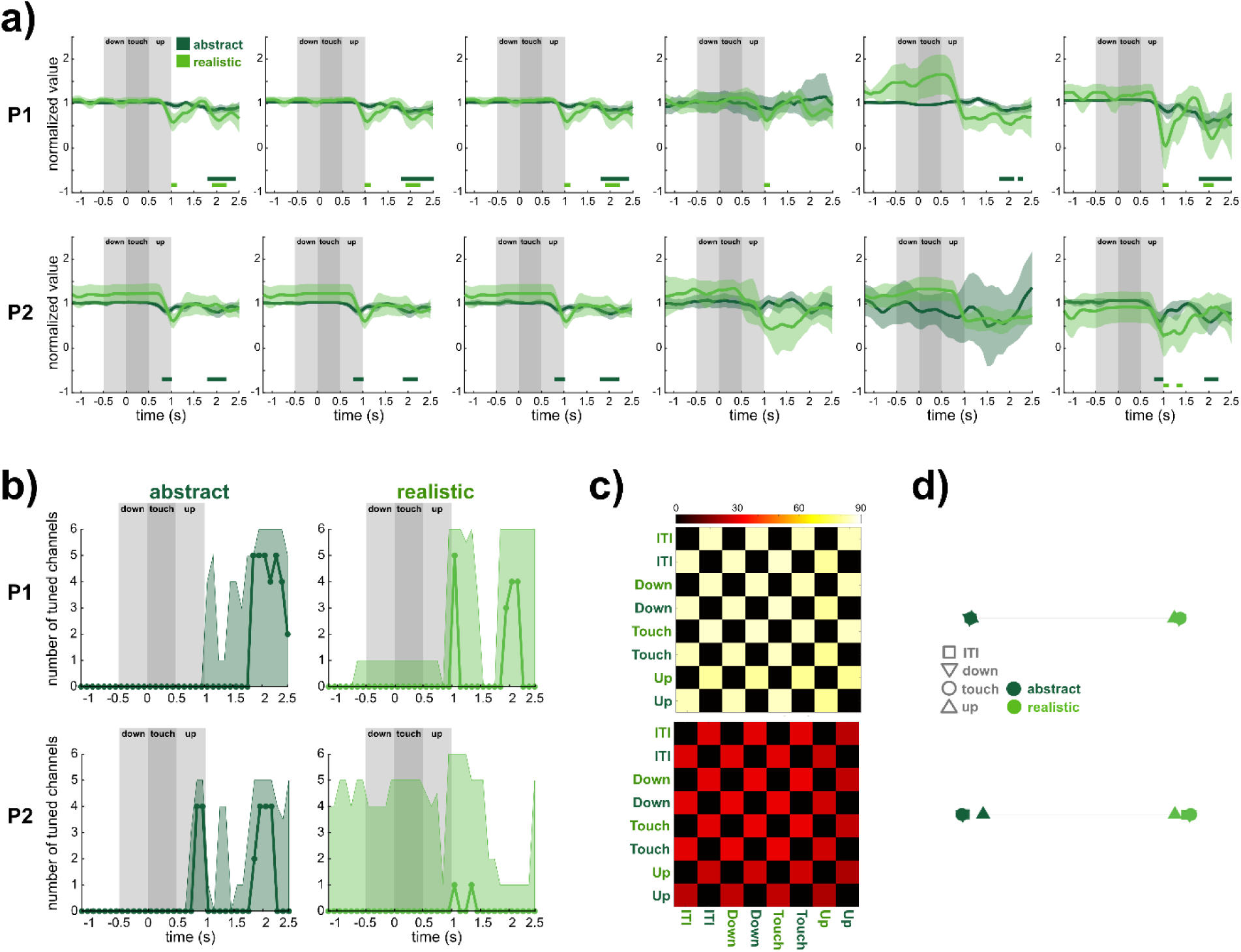
Eye movements during catch trials. **a)** 6 eye movement features were recorded in each participant. The average value of each feature is plotted, averaged across catch trials, by condition. Left-most 3 plots represent the coordinates used to encode eye position within the 3D virtual environment, and right-most 3 plots represent the vector of eye gaze direction. Analysis, plotting follow methods of Fig. 5b. **b)** Percentage of features (n=6) tuned during catch trials relative to baseline, separated by condition. Time is aligned to the onset of the “touch” phase of the visual stimulus. Shaded area indicates 95% CIs computed by bootstrapping across trials (P1 n=12 realistic, 36 abstract trials, P2 n=26 realistic, 30 abstract trials) over 1000 iterations. Analysis, plotting follow methods of Fig. 5a. **c)** Representational dissimilarity matrices (RDM) of eye movement features Analysis, plotting follow methods of Fig. 5d. **d)** Multi-dimensional scaling (MDS) plots of the RDMs in **(c)**. Grey lines between icons are “rubber bands” whose thickness is based on the goodness of fit of the scaling (P1: Pearson’s r=0.99, p=2.3x10^-46^; P2: r=0.99, p=2.5x10^-31^). Analysis, plotting follow methods of Fig. 5e.

## References

1. Ariani, G., Pruszynski, J.A., Diedrichsen, J., 2022. Motor planning brings human primary somatosensory cortex into action-specific preparatory states. eLife 11, e69517. 10.7554/eLife.69517

2. Armenta Salas, M., Bashford, L., Kellis, S., Jafari, M., Jo, H., Kramer, D., Shanfield, K., Pejsa, K., Lee, B., Liu, C.Y., Andersen, R.A., 2018. Proprioceptive and cutaneous sensations in humans elicited by intracortical microstimulation. eLife 7. 10.7554/eLife.32904

3. Bashford, L., Rosenthal, I., Kellis, S., Pejsa, K., Kramer, D., Lee, B., Liu, C., Andersen, R.A., 2021. The neurophysiological representation of imagined somatosensory percepts in human cortex. J. Neurosci. 41, 2177–2185. 10.1523/JNEUROSCI.2460-20.2021

4. Berg, J.A., Dammann, III, J.F., Tenore, F.V., Tabot, G.A., Boback, J.L., Manfredi, L.R., Peterson, M.L., Katyal, K.D., Johannes, M.S., Makhlin, A., Wilcox, R., Franklin, R.K., Vogelstein, R.J., Hatsopoulos, N.G., Bensmaia, S.J., 2013. Behavioral demonstration of a somatosensory neuroprosthesis. IEEE Trans. Neural Syst. Rehabil. Eng. 21, 500–507. 10.1109/TNSRE.2013.2244616

5. Bjånes, D.A., Bashford, L., Pejsa, K., Lee, B., Liu, C.Y., Andersen, R.A., 2022. Multi-channel intra-cortical micro-stimulation yields quick reaction times and evokes natural somatosensations in a human participant (preprint). Medarxiv. 10.1101/2022.08.08.22278389

6. Blakemore, S.-J., Bristow, D., Bird, G., Frith, C., Ward, J., 2005. Somatosensory activations during the observation of touch and a case of vision–touch synaesthesia. Brain 128, 1571–1583. 10.1093/brain/awh500

7. Bufalari, I., Aprile, T., Avenanti, A., Di Russo, F., Aglioti, S.M., 2007. Empathy for pain and touch in the human somatosensory cortex. Cereb. Cortex 17, 2553–2561. 10.1093/cercor/bhl161

8. Caldwell, D.J., Cronin, J.A., Wu, J., Weaver, K.E., Ko, A.L., Rao, R.P.N., Ojemann, J.G., 2019. Direct stimulation of somatosensory cortex results in slower reaction times compared to peripheral touch in humans. Sci. Rep. 9. 10.1038/s41598-019-38619-2

9. Callier, T., Brantly, N.W., Caravelli, A., Bensmaia, S.J., 2020. The frequency of cortical microstimulation shapes artificial touch. Proc. Natl. Acad. Sci. U. S. A. 117, 1191–1200. 10.1073/pnas.1916453117

10. Chan, A.W.-Y., Baker, C.I., 2015. Seeing is not feeling: Posterior parietal but not somatosensory cortex engagement during touch observation. J. Neurosci. 35, 1468–1480. 10.1523/JNEUROSCI.3621-14.2015

11. Chapman, C.E., Meftah, E.-M., 2005. Independent controls of attentional influences in primary and secondary somatosensory cortex. J. Neurophysiol. 94, 4094–4107. 10.1152/jn.00303.2005

12. Christie, B.P., Charkhkar, H., Shell, C.E., Marasco, P.D., Tyler, D.J., Triolo, R.J., 2019a. Visual inputs and postural manipulations affect the location of somatosensory percepts elicited by electrical stimulation. Sci. Rep. 9, 11699. 10.1038/s41598-019-47867-1

13. Christie, B.P., Graczyk, E.L., Charkhkar, H., Tyler, D.J., Triolo, R.J., 2019b. Visuotactile synchrony of stimulation-induced sensation and natural somatosensation. J. Neural Eng. 16, 036025. 10.1088/1741-2552/ab154c

14. Christie, B.P., Osborn, L.E., McMullen, D.P., Pawar, A.S., Thomas, T.M., Bensmaia, S.J., Celnik, P.A., Fifer, M.S., Tenore, F.V., 2022. Perceived timing of cutaneous vibration and intracortical microstimulation of human somatosensory cortex. Brain Stimulat. 15, 881–888. 10.1016/j.brs.2022.05.015

15. Christie, B.P., Tat, D.M., Irwin, Z.T., Gilja, V., Nuyujukian, P., Foster, J.D., Ryu, S.I., Shenoy, K.V., Thompson, D.E., Chestek, C.A., 2015. Comparison of spike sorting and thresholding of voltage waveforms for intracortical brain-machine interface performance. J. Neural Eng. 12, 016009. 10.1088/1741-2560/12/1/016009

16. Colino, F.L., Lee, J.-H., Binsted, G., 2017. Availability of vision and tactile gating: vision enhances tactile sensitivity. Exp. Brain Res. 235, 341–348. 10.1007/s00221-016-4785-3

17. Collinger, J.L., Wodlinger, B., Downey, J.E., Wang, W., Tyler-Kabara, E.C., Weber, D.J., McMorland, A.J., Velliste, M., Boninger, M.L., Schwartz, A.B., 2013. High-performance neuroprosthetic control by an individual with tetraplegia. The Lancet 381, 557–564. 10.1016/S0140-6736(12)61816-9

18. Dai, J., Zhang, P., Sun, H., Qiao, X., Zhao, Y., Ma, J., Li, S., Zhou, J., Wang, C., 2019. Reliability of motor and sensory neural decoding by threshold crossings for intracortical brain–machine interface. J. Neural Eng. 16, 036011. 10.1088/1741-2552/ab0bfb

19. Dekleva, B.M., Weiss, J.M., Boninger, M.L., Collinger, J.L., 2021. Generalizable cursor click decoding using grasp-related neural transients. J. Neural Eng. 18, 10.1088/1741-2552/ac16b2. https://doi.org/10.1088/1741-2552/ac16b2

20. Diedrichsen, J., Berlot, E., Mur, M., Schütt, H.H., Shahbazi, M., Kriegeskorte, N., 2021. Comparing representational geometries using whitened unbiased-distance-matrix similarity. Neurons Behav. Data Anal. Theory 5, 1–31. 10.51628/001c.27664

21. Dionne, J.K., Legon, W., Staines, W.R., 2013. Crossmodal influences on early somatosensory processing: interaction of vision, touch, and task-relevance. Exp. Brain Res. 226, 503–512. 10.1007/s00221-013-3462-z

22. Ebisch, S.J.H., Perrucci, M.G., Ferretti, A., Del Gratta, C., Romani, G.L., Gallese, V., 2008. The sense of touch: Embodied simulation in a visuotactile mirroring mechanism for observed animate or inanimate touch. J. Cogn. Neurosci. 20, 1611–1623. 10.1162/jocn.2008.20111

23. Espenhahn, S., Yan, T., Beltrano, W., Kaur, S., Godfrey, K., Cortese, F., Bray, S., Harris, A.D., 2020. The effect of movie-watching on electroencephalographic responses to tactile stimulation. NeuroImage 220, 117130. 10.1016/j.neuroimage.2020.117130

24. Flesher, S.N., Collinger, J.L., Foldes, S.T., Weiss, J.M., Downey, J.E., Tyler-Kabara, E.C., Bensmaia, S.J., Schwartz, A.B., Boninger, M.L., Gaunt, R.A., 2016. Intracortical microstimulation of human somatosensory cortex. Sci. Transl. Med. 8, 361ra141. 10.1126/scitranslmed.aaf8083

25. Flesher, S.N., Downey, J.E., Weiss, J.M., Hughes, C.L., Herrera, A.J., Tyler-Kabara, E.C., Boninger, M.L., Collinger, J.L., Gaunt, R.A., 2021. A brain-computer interface that evokes tactile sensations improves robotic arm control. Science 372, 831–836. 10.1126/science.abd0380

26. Gale, D.J., Flanagan, J.R., Gallivan, J.P., 2021. Human somatosensory cortex is modulated during motor planning. J. Neurosci. 41, 5909–5922. 10.1523/JNEUROSCI.0342-21.2021

27. Ghez, C., Gordon, J., Ghilardi, M.F., 1995. Impairments of reaching movements in patients without proprioception. II. Effects of visual information on accuracy. J. Neurophysiol. 73, 361–372. 10.1152/jn.1995.73.1.361

28. Giummarra, M.J., Gibson, S.J., Georgiou-Karistianis, N., Bradshaw, J.L., 2008. Mechanisms underlying embodiment, disembodiment and loss of embodiment. Neurosci. Biobehav. Rev. 32, 143–160. 10.1016/j.neubiorev.2007.07.001

29. Godlove, J.M., Whaite, E.O., Batista, A.P., 2014. Comparing temporal aspects of visual, tactile, and microstimulation feedback for motor control. J. Neural Eng. 11, 046025. 10.1088/1741-2560/11/4/046025

30. Gonzalez-Franco, M., Ofek, E., Pan, Y., Antley, A., Steed, A., Spanlang, B., Maselli, A., Banakou, D., Pelechano, N., Orts-Escolano, S., Orvalho, V., Trutoiu, L., Wojcik, M., Sanchez-Vives, M.V., Bailenson, J., Slater, M., Lanier, J., 2020. The Rocketbox library and the utility of freely available rigged avatars. Front. Virtual Real. 1, 561558. 10.3389/frvir.2020.561558

31. Grunewald, A., Linden, J.F., Andersen, R.A., 1999. Responses to Auditory Stimuli in Macaque Lateral Intraparietal Area I. Effects of Training. J. Neurophysiol. 82, 330–342. 10.1152/jn.1999.82.1.330

32. Haggard, P., Christakou, A., Serino, A., 2007. Viewing the body modulates tactile receptive fields. Exp. Brain Res. 180, 187–193. 10.1007/s00221-007-0971-7

33. Hughes, C.L., Flesher, S.N., Gaunt, R.A., 2022. Effects of stimulus pulse rate on somatosensory adaptation in the human cortex. Brain Stimulat. 15, 987–995. 10.1016/j.brs.2022.05.021

34. Hughes, C.L., Flesher, S.N., Weiss, J.M., Boninger, M.L., Collinger, J., Gaunt, R., 2021. Perception of microstimulation frequency in human somatosensory cortex. eLife 10, e65128. 10.7554/eLife.65128

35. Jafari, M., Aflalo, T., Chivukula, S., Kellis, S.S., Salas, M.A., Norman, S.L., Pejsa, K., Liu, C.Y., Andersen, R.A., 2020. The human primary somatosensory cortex encodes imagined movement in the absence of sensory information. Commun. Biol. 3, 757. 10.1038/s42003-020-01484-1

36. Jeannerod, M., 2003. The mechanism of self-recognition in humans. Behav. Brain Res. 142, 1–15. 10.1016/S0166-4328(02)00384-4

37. Kennett, S., Taylor-Clarke, M., Haggard, P., 2001. Noninformative vision improves the spatial resolution of touch in humans. Curr. Biol. 11, 1188–1191. 10.1016/S0960-9822(01)00327-X

38. Keshtkaran, M.R., Sedler, A.R., Chowdhury, R.H., Tandon, R., Basrai, D., Nguyen, S.L., Sohn, H., Jazayeri, M., Miller, L.E., Pandarinath, C., 2022. A large-scale neural network training framework for generalized estimation of single-trial population dynamics. Nat. Methods 19, 1572–1577. 10.1038/s41592-022-01675-0

39. Keysers, C., Wicker, B., Gazzola, V., Anton, J.-L., Fogassi, L., Gallese, V., 2004. A touching sight: SII/PV activation during the observation and experience of touch. Neuron 42, 335–346. 10.1016/S0896-6273(04)00156-4

40. Kimura, T., 2021. Approach of visual stimuli facilitates the prediction of tactile events and suppresses beta band oscillations around the primary somatosensory area. Neuroreport 32, 631–635. 10.1097/WNR.0000000000001643

41. Klaes, C., Shi, Y., Kellis, S., Minxha, J., Revechkis, B., Andersen, R.A., 2014. A cognitive neuroprosthetic that uses cortical stimulation for somatosensory feedback. J. Neural Eng. 11. 10.1088/1741-2560/11/5/056024

42. Kriegeskorte, N., 2008. Representational similarity analysis – connecting the branches of systems neuroscience. Front. Syst. Neurosci. 2. 10.3389/neuro.06.004.2008

43. Kuehn, E., Haggard, P., Villringer, A., Pleger, B., Sereno, M.I., 2018. Visually-driven maps in Area 3b. J. Neurosci. 38, 1295–1310. 10.1523/JNEUROSCI.0491-17.2017

44. Kuehn, E., Trampel, R., Mueller, K., Turner, R., Schütz-Bosbach, S., 2013. Judging roughness by sight-A 7- tesla fMRI study on responsivity of the primary somatosensory cortex during observed touch of self and others. Hum. Brain Mapp. 34, 1882–1895. 10.1002/hbm.22031

45. Lafer-Sousa, R., Hermann, K.L., Conway, B.R., 2015. Striking individual differences in color perception uncovered by ‘the dress’ photograph. Curr. Biol. 25, R545–R546. 10.1016/j.cub.2015.04.053

46. Lee, J., Maunsell, J.H.R., 2009. A Normalization Model of Attentional Modulation of Single Unit Responses. PLoS ONE 4, e4651. 10.1371/journal.pone.0004651

47. Linden, J.F., Grunewald, A., Andersen, R.A., 1999. Responses to Auditory Stimuli in Macaque Lateral Intraparietal Area II. Behavioral Modulation. J. Neurophysiol. 82, 343–358. 10.1152/jn.1999.82.1.343

48. Longo, M.R., Pernigo, S., Haggard, P., 2011. Vision of the body modulates processing in primary somatosensory cortex. Neurosci. Lett. 489, 159–163. 10.1016/j.neulet.2010.12.007

49. Miall, R.C., Afanasyeva, D., Cole, J.D., Mason, P., 2021. The role of somatosensation in automatic visuo- motor control: a comparison of congenital and acquired sensory loss. Exp. Brain Res. 239, 2043– 2061. 10.1007/s00221-021-06110-y

50. Miall, R.C., Rosenthal, O., Ørstavik, K., Cole, J.D., Sarlegna, F.R., 2019. Loss of haptic feedback impairs control of hand posture: a study in chronically deafferented individuals when grasping and lifting objects. Exp. Brain Res. 237, 2167–2184. 10.1007/s00221-019-05583-2

51. Morrison, I., Lloyd, D., Di Pellegrino, G., Roberts, N., 2004. Vicarious responses to pain in anterior cingulate cortex: Is empathy a multisensory issue? Cogn. Affect. Behav. Neurosci. 4, 270–278. 10.3758/CABN.4.2.270

52. Moses, D.A., Metzger, S.L., Liu, J.R., Anumanchipalli, G.K., Makin, J.G., Sun, P.F., Chartier, J., Dougherty, M.E., Liu, P.M., Abrams, G.M., Tu-Chan, A., Ganguly, K., Chang, E.F., 2021. Neuroprosthesis for decoding speech in a paralyzed person with anarthria. N. Engl. J. Med. 385, 217–227. 10.1056/NEJMoa2027540

53. Nili, H., Wingfield, C., Walther, A., Su, L., Marslen-Wilson, W., Kriegeskorte, N., 2014. A toolbox for representational similarity analysis. PLoS Comput. Biol. 10, e1003553. 10.1371/journal.pcbi.1003553

54. Pandarinath, C., Bensmaia, S.J., 2022. The science and engineering behind sensitized brain-controlled bionic hands. Physiol. Rev. 102, 551–604. 10.1152/physrev.00034.2020

55. Popovich, C., Staines, W.R., 2014. The attentional-relevance and temporal dynamics of visual-tactile crossmodal interactions differentially influence early stages of somatosensory processing. Brain Behav. 4, 247–260. 10.1002/brb3.210

56. Press, C., Taylor-Clarke, M., Kennett, S., Haggard, P., 2004. Visual enhancement of touch in spatial body representation. Exp. Brain Res. 154, 238–245. 10.1007/s00221-003-1651-x

57. Reynolds, J.H., Heeger, D.J., 2009. The Normalization Model of Attention. Neuron 61, 168–185. 10.1016/j.neuron.2009.01.002

58. Risso, G., Valle, G., 2022. Multisensory integration in bionics: Relevance and perspectives. Curr. Phys. Med. Rehabil. Rep. 10.1007/s40141-022-00350-x

59. Robles-De-La-Torre, G., 2006. The importance of the sense of touch in virtual and real environments. IEEE Multimed. 13, 24–30. 10.1109/MMUL.2006.69

60. Rosenthal, I.A., Bashford, L., Kellis, S., Pejsa, K., Lee, B., Liu, C., Andersen, R.A., 2023. S1 represents multisensory contexts and somatotopic locations within and outside the bounds of the cortical homunculus. Cell Rep. 42, 112312. 10.1016/j.celrep.2023.112312

61. Sainburg, R.L., Ghilardi, M.F., Poizner, H., Ghez, C., 1995. Control of limb dynamics in normal subjects and patients without proprioception. J. Neurophysiol. 73, 820–835. 10.1152/jn.1995.73.2.820

62. Schaefer, M., Xu, B., Flor, H., Cohen, L.G., 2009. Effects of different viewing perspectives on somatosensory activations during observation of touch. Hum. Brain Mapp. 30, 2722–2730. 10.1002/hbm.20701

63. Simeral, J.D., Hosman, T., Saab, J., Flesher, S.N., Vilela, M., Franco, B., Kelemen, J., Brandman, D.M., Ciancibello, J.G., Rezaii, P.G., Eskandar, E.N., Rosler, D.M., Shenoy, K.V., Henderson, J.M., Nurmikko, A.V., Hochberg, L.R., 2021. Home use of a percutaneous wireless intracortical brain-computer interface by individuals with tetraplegia. IEEE Trans. Biomed. Eng. 1–1. 10.1109/TBME.2021.3069119

64. Tabot, G.A., Kim, S.S., Winberry, J.E., Bensmaia, S.J., 2015. Restoring tactile and proprioceptive sensation through a brain interface. Neurobiol. Dis. 83, 191–198. 10.1016/j.nbd.2014.08.029

65. Tipper, S., Phillips, N., Dancer, C., Lloyd, D., Howard, L., McGlone, F., 2001. Vision influences tactile perception at body sites that cannot be viewed directly. Exp. Brain Res. 139, 160–167. 10.1007/s002210100743

66. Tsakiris, M., Carpenter, L., James, D., Fotopoulou, A., 2010. Hands only illusion: multisensory integration elicits sense of ownership for body parts but not for non-corporeal objects. Exp. Brain Res. 204, 343–352. 10.1007/s00221-009-2039-3

67. Walther, A., Nili, H., Ejaz, N., Alink, A., Kriegeskorte, N., Diedrichsen, J., 2016. Reliability of dissimilarity measures for multi-voxel pattern analysis. NeuroImage 137, 188–200. 10.1016/j.neuroimage.2015.12.012

68. Willsey, M.S., Nason-Tomaszewski, S.R., Ensel, S.R., Temmar, H., Mender, M.J., Costello, J.T., Patil, P.G., Chestek, C.A., 2022. Real-time brain-machine interface in non-human primates achieves high-velocity prosthetic finger movements using a shallow feedforward neural network decoder. Nat. Commun. 13, 6899. 10.1038/s41467-022-34452-w

69. Yoo, S.-S., Freeman, D.K., McCarthy, J.J., Jolesz, F.A., 2003. Neural substrates of tactile imagery: a functional MRI study. NeuroReport 14, 581–585. 10.1097/00001756-200303240-00011

