## Supplementary figures and images for "Visual context affects the perceived timing of tactile sensations elicited through intra-cortical microstimulation"

### Supplementary Figure 1

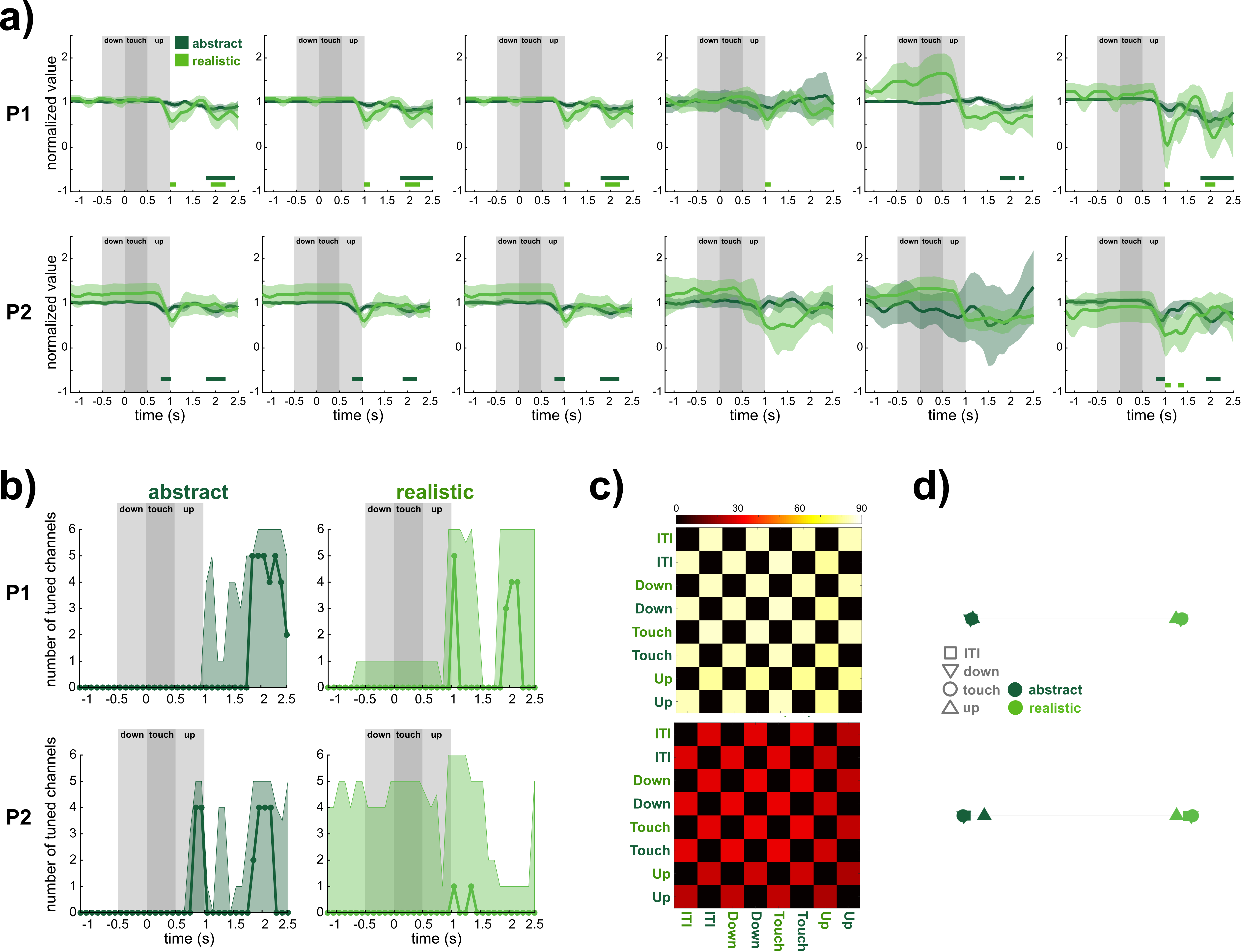
